# Nanofibrous Wound Dressing with a Smart Drug Delivery System: Poly(N-Isopropylacrylamide)-Conjugated Polycaprolactone Nanofibers Loaded with Curcumin

**DOI:** 10.1101/2024.06.04.597437

**Authors:** Dawood Khan, Anirban Jyoti, Nik Aliaa, Kapil D. Patel, Renjian Tan, Hae-Won Kim, David Chau, Linh Nguyen

## Abstract

In response to clinical demands for advanced wound dressings, a "smart bandage" was developed by combining polycaprolactone (PCL) nanofibers and the thermoresponsive polymer poly(N-isopropylacrylamide) (PNIPAAm). This bandage incorporates curcumin, an anti-inflammatory and antioxidant agent, to enhance wound healing. The LCST of PNIPAAm (32°C) enables an "on-off" drug delivery system, transitioning from a hydrophilic coil to a hydrophobic state. The nanofiber matrix was characterized using FTIR spectroscopy, SEM, water contact angle, and dynamic mechanical analysis. Curcumin release was assessed both above and below the LCST. The bandage is comfortable, easy to apply and remove, and promotes rapid wound healing. In vitro studies confirmed its non-toxicity to human dermal fibroblast cells. This "smart bandage" represents a significant advancement in wound care, with the potential to meet clinical requirements and enhance patient compliance, offering controlled drug delivery combined with the therapeutic benefits of curcumin.

## 1. Introduction

Wounds are disruptions of the normal structure and function of the skin and underlying tissues, which can result from surgery, trauma, or disease [1,2]. This definition encompasses a wide range of injuries that vary in severity, location, and cause. The skin is the largest organ of the body and serves as a protective barrier against external threats, such as infection, dehydration, and mechanical damage [3–5]. The underlying tissues, such as muscles, tendons, nerves, and blood vessels, provide support, movement, sensation, and nutrition to the skin and other organs. Therefore, any wound that affects the skin and/or the underlying tissues can have significant consequences for the health and well-being of the individual [6,7]. Thus, wounds can cause significant pain and discomfort to the patients, as well as pose challenges for wound management and infection prevention. Wounds can also lead to serious complications, such as loss of function, tissue necrosis, amputation, or even death [8]. Wounds require prompt medical attention, effective treatment, and skilled care to prevent potential complications, such as infection, scarring, or chronicity [8, 9,10].

Wound healing is a vital biological process that restores the integrity and functionality of the skin and underlying tissues after injury or disease [1–10]. However, wound healing can be impaired or delayed by various intrinsic and extrinsic factors, such as oxygenation, infection, inflammation, diabetes, vascular disease, or malnutrition [9–11]. Depending on the duration and progress of healing, wounds can be classified as acute or chronic. Acute wounds heal in a timely and orderly manner, following four phases: hemostasis, inflammation, proliferation, and maturation. Chronic wounds, on the other hand, remain in a prolonged inflammatory state and fail to heal properly [8]. Chronic wounds are more prevalent and problematic than acute wounds, affecting millions of people worldwide and imposing a significant burden on the health care system [12]. In the UK alone, there were an estimated 3.8 million patients with a wound managed by the NHS in 2017/2018, of which 70% healed in the study year. The annual NHS cost of wound management was £8.3 billion, of which £2.7 billion and £5.6 billion were associated with managing healed and unhealed wounds, respectively [13]. Therefore, it is essential to develop effective treatments that can enhance wound healing and prevent complications. One of the promising strategies is to use biomaterials that can interact with the wound environment and promote tissue regeneration. Among these biomaterials, smart biomaterials are those that can sense and respond to external stimuli or changes in the wound condition, such as temperature, pH, or enzyme activity [14]. Although, smart biomaterials can offer advantages such as controlled drug delivery, adaptive degradation or swelling, and wound status monitoring, numerous barriers hinder the development of wound healing bandages, including a limited understanding of cellular mechanisms and tissue regeneration, as well as a lack of preclinical and clinical trials, which restrict the exploration, potential, and scalability of various products [14, 15]. Nonetheless, numerous studies have been conducted and reported with a primary focus on enhancing the healing rate. These studies have explored novel material formulations, drug delivery systems, and tissue replacements [16–18]. Patient compliance plays a crucial role in effective wound management, where ease of application and removal are key considerations. Prioritizing patient compliance not only enhances their quality of life but also eases the burden on clinical settings. Therefore, the development of advanced and functional wound dressings can substantially improve patients’ lifestyles and provide a foundation for accelerated wound recovery and regeneration.

Existing therapeutic options encompass a wide spectrum, ranging from biological and synthetic dressings to those incorporating pharmaceutical agents [19–22]. Although, they are designed to promote wound healing and tissue regeneration, however each of them has their own advantages and disadvantages. For example, biological dressings, encompassing hyaluronic acid, chitosan, collagen, elastin, and alginate, have garnered significant attention due to their inherent sustainability [22–25]. However, challenges arise from their variability in properties, limited scalability in production, and impurity concerns. Consequently, a common strategy is to combine these biological dressings with synthetic materials to maximize their full potential and enhance their [22–25]. In contrast, synthetic dressings, including hydrogels, foams, hydrocolloids, films, and nanofibers, offer an even higher degree of controllability over their properties, which holds the potential for improving moisture retention, neovascularization, growth factor delivery, tissue adhesion, and antimicrobial properties within the wound environment [26]. Nevertheless, the effectiveness of these synthetic dressings depends on the wound’s severity and condition. For example, hydrogel dressings like polyvinyl pyrrolidone (PVP) and poly (methyl methacrylate) (PMMA), with their high water content and flexibility facilitating efficient metabolite diffusion, are still challenged by mechanical stability and wound exudate accumulation issues, which can significantly affect patient compliance [22, 27].

Nanofibers represent one of the most promising forms of wound dressings, addressing several limitations observed in other dressing materials. In this study, we have employed electrospinning, a scalable, cost-effective, and versatile technique [28]. This technique enables the fabrication of composites with high and tunable porosity, a high surface area-to-volume ratio, and the capacity to produce various shapes and sizes, thereby enhancing the final properties and functionalities of the dressings [29].

In the current study, a well-studied thermo-responsive polymer was used of poly(N-isopropylacrylamide) (PNIPAAm), to exploit its thermal transition property [30]. This unique behaviour allows a controlled as well as an efficient temperature-dependent drug delivery system. Furthermore, the incorporation of surfactants, salts, or copolymerization has been extensively investigated for surface functionalization, enabling the fine-tuning of the LCST of PNIPAAm and expanding its potential applications [31]. Curcumin is a common natural herbal medicine that holds potential in facilitating wound healing, as it is known to possess antioxidant and polyphenolic compounds which offers free-radical scavenging and anti-inflammatory activities, that assist in preventing oxidative stress, in which over-exposure of the stress could further damage the surrounding cells and delayed the rate of wound healing [32–34]. Curcumin also known to facilitate formation of granulation tissue and blood vessels, re-epithelialization, deposition of high collagen content (type III relative to type I) and attracting vital cells to the wound site during proliferation stage [35–36]. However, its hydrophobic property leads to some drawbacks such as mostly insoluble in aqueous media, short half-life and fast metabolism, and the polyphenol moiety compound within curcumin raise a toxic response which needs to be addressed to achieve better solubility, stability and safe release for drug delivery system [36–37]. Hence to address barriers and facilitate advancement in the formulation of wound dressings, this project aims to produce a novel PCL nanofiber grafted with PNIPAAm through amidation process and loaded with curcumin to produce the overall matrix PCL-PNIPAAm-Cur via electrospinning.

## 2. Materials and methods

### 2.1 Materials

Sodium hydroxide (NaOH, 98%), MES hydrate (99.5%), N-Hydroxysuccinimide (NHS, 98%), N-(3-Dimethylaminopropyl)-N’-ethylcarbodiimide hydrochloride (EDC, crystalline) (E7750), Poly(N-isopropylacrylamide), amine terminated, average Mn 2,500 (T), penicillin/streptomycin (P/S) and curcumin were purchased from Sigma-Aldrich, UK.

Dulbecco’s Modified Eagle Medium (DMEM), foetal bovine serum (FBS), phosphate-buffered saline (PBS, pH 7.2), LIVE/DEAD Viability/Cytotoxicity Kit, alamarBlue^TM^ Cell Viability Reagent and human dermal fibroblast (HDF) cells (C0135C) were obtained from ThermoFisher Scientific, UK. Ethanol (95+%, Acros Organics^TM^, 10318820) was purchased from Fisher Scientific, UK.

### 2.2 Fabrication of PCL nanofibres

PCL (MW = 80,000, Sigma-Aldrich) solution (13% w/v) was prepared in a mixture of chloroform (CHL) and tetrafluoroethanol (TFE) solvent at a CHL:TFE ratio of 4:1. The electrospinning parameters were optimised at applied voltage (13kV), flow rate (0.5 ml/h), needle size (21-G), tip-to-collector distance (22-25 cm) and collector speed (200 rpm) to produce these nanofibres. This was performed according to the methodology described by Patel et al., (25).

### 2.3 Preparation of PCL-PNIPAAm-Cur membranes

PCL nanofibrous sheets were cut into discs (16 mm diameter, n=24) using an Acuderm Acu-Punch Biopsy Punch® (Acuderm Inc., Fisher Scientific Ltd., Loughborough, UK) and inserted into a 24-well plate. A measured volume of NaOH solution (20 ml, 1 M) was then added equally into each plate using a micropipette and stirred using a rocker shaker (90 rpm) for 3 hours. Subsequently, an activation solution was prepared by adding MES hydrate (0.195 g, 0.05 M), NHS (0.138 g, 0.06 M) and EDC (0.460 g, 0.12 M) into deionised water (20 ml). The pH of the activation solution was adjusted and confirmed to be 6 using pH indicator strips (Whatman International Ltd., Maidstone, UK). The NaOH solution was then removed, and the membranes were washed once with deionised water. The activation solution was then added to the PCL membranes and stirred again (90 rpm) for 3 hours to activate them for surface modification. Following this, the activation solution was removed and stored in another beaker. Amine terminated PNIPAAm (1 g) was added and dissolved in the activation solution. The PCL membranes were then reintroduced to the new activation solution and stirred (90 rpm) at 4 °C for 24 hours. This was performed according to the methodology described by Nguyen et al. (26). The new activation solution was then removed, and the membranes (PCL-PNIPAAm) were dried in a fume hood for 3 hours and stored at 4 °C overnight. They were then washed once with deionised water and 200 µl of curcumin (1mg/ml in 50% ethanol solution) was then added to each well containing the membranes and stored at 4 °C overnight.

### 2.4 Characterisation of PCL, PCL-PNIPAAm and PCL-PNIPAAm-Cur

#### 2.4.1 Lower critical solution temperature (LCST) measurements

The lower critical solution temperature (LCST) transition properties of the polymer solutions were investigated by UV-visible measurements at different temperatures (ranging from 20°C to 40°C). 1 g of PNIPAAm (Product Number: 724823) was added to 10 ml of distilled water to prepare polymer solution. 100 µl of the solution were added into a 96 well plate (CLS3635) (Corning UV-Transparent Microplates). The absorbance of the solution was measured by microplate reader (Tecan Infinite M200, Tecan Ltd., Reading, UK) from Tecan at 563 nm wavelength with temperature increased from 20°C to 40°C. The cloud point temperature of the solution is determined by when the absorbance is raised to 50% of the initial value.

#### 2.4.2 Chemical composition analysis

Functional chemical groups of PCL, PCL-PNIPAAm, and PCL-PNIPAAm-Cur membranes were characterised using a attenuated total reflectance Fourier transform infrared (ATR-FTIR) spectrophotometer (Spectrum One, PerkinElmer, Llanstrisant, UK). The software Time Base (Spectrum) was used to process the spectra, whereby the samples were scanned in the range of 800 – 4000 cm^-1^. A blank background absorbance was recorded first before taking readings of the samples.

#### 2.4.3 Morphology analysis

Surface morphology and size distribution of the fibres were determined using scanning electron microscopy (SEM; Zeiss EVO HD, Jena, Germany). The PCL, PCL-PNIPAAm, and PCL-PNIPAAm-Cur membranes were cut into discs (16mm diameter, n=2) and then coated with 95% gold and 5% palladium (Polaron E5000 Sputter Coater, Quorum Technologies, UK). The electrospun membranes were observed at both 1000X and 5000X magnification.

#### 2.4.4 Contact angle measurement

Hydrophilicity/hydrophobicity of the electrospun PCL, PCL-PNIPAAm, and PCL-PNIPAAm-Cur membranes were measured and compared using a Contact Angle Measuring System (KSV CAM200 Optical Contact Angle Meter, Biolin Scientific, UK) at room temperature (19 °C). The membranes were placed on a sample stage, and a water droplet was dropped automatically onto the surface of the sample to gage the water contact angle. A thin needle was used to make sessile drops. The drop was illuminated from one side and the image of the water drop was recorded by a camera at the opposite side of the instrument. The drop image was transferred to a computer and shown on the monitor. The CAM software analysed the drop image to calculate the contact angle.

#### 2.4.5 Tensile strength measurement

Rectangular samples (∼5 x 20 mm, n=5) of the PCL and PCL-PNIPAAm membranes were cut and tensile strength tests were performed using Dynamic Mechanical Analysis (DMA 850; TA Instruments, New Castle, USA) at room temperature. The ultimate tensile strength, % elongation and Young’s modulus were determined using TRIOS software.

#### 2.4.6 Drug-release profiles

Drug-release profiles of curcumin were determined from PCL-Cur and PCL-PNIPAAM-Cur membranes at different temperatures. They were each cut into discs (16 mm diameter, n=5) and immersed in ethanol (50%, 1.5 ml) in a 24-well plate at 4 °C. The samples were also set up at 37 °C as a control to compare release profiles above the LCST of PNIPAAm (32 °C). The ethanol was removed from the solutions and analysed at regular intervals (1, 2, 3, 4, 5 hours) to measure the ‘initial burst’ release profile. Further analysis was carried out at 1 and 5 days to determine the ‘absolute’ release profile. The samples (100 µl) were decanted into a 96 well-plate (CLS3635; Corning UV-Transparent Microplates) and analysed using UV absorbance from a microplate reader (Tecan Infinite M200, Tecan Ltd., Reading, UK) between 350-500 nm. The concentration of curcumin release was quantified from a calibration curve of known concentrations. All the samples were diluted 10 times and the formulae used for calculations to obtain the cumulative percentage release of curcumin from all the membranes are as follows:

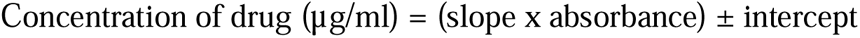

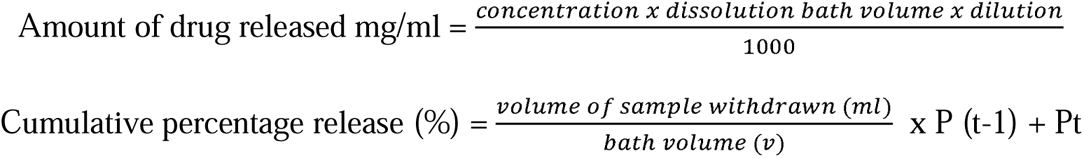

Where Pt = Percentage release at time t

Where P (t-1) = Percentage release before ‘t’

This was performed according to the methodology described by Chandrasekaran et al., (27). Microsoft Excel 2019 was used to analyse the data.

### 2.5 In-vitro study

#### 2.5.1 Cell culture

Human dermal fibroblast (HDF) cells (C0135C; ThermoFisher Scientific, UK) were cultured in flasks with Dulbecco’s Modified Eagle Medium (DMEM; ThermoFisher Scientific, UK) supplemented with 10% foetal bovine serum (FBS) and 100 U/ml of penicillin and 100 µg/ml of streptomycin (PS) under standard cell culture conditions (37 °C, 5% CO_2_, 95% humidity). The cells were seeded on the electrospun membranes at a density of 10^4^ cells/ml/sample in a 24-well plate with culture medium.

#### 2.5.2 Cell viability and metabolic activity assays

For cytotoxicity, cell adhesion and cell growth assessment of the nanofibrous membranes, HDF cells were employed as an appropriate *in vitro* system. The PCL, PCL-PNIPAAm and PCL-PNIPAAm-Cur membranes were sterilized (UV, Steristorm 2537a, Coast-Air, London, UK) for 20 minutes and soaked in PBS for 1 minute before seeding. The HDF cells were then cultured on these samples and evaluated after 1, 3, and 7 days of incubation under standard cell culture conditions using the LIVE/DEAD® Viability/Cytotoxicity Kit and alamarBlue^TM^ Cell Viability Reagent. In addition, HDF cells in media, without the presence of a membrane, was used as a control.

LIVE/DEAD staining is a dye-based assay that can only give a direct viability assessment, whereby cells are either alive (green) or dead (red). HDF cells were seeded onto the control and discs of PCL, PCL-PNIPAAm, PCL-PNIPAAm-Cur membranes (16 mm diameter, n=2) in 24-well plates at a density of 1 x 10^4^ cells/ml. 1 ml of media was added into each well. The media was discarded, and the samples were washed with PBS at each timepoint. A stock solution of EthD-1 (20 µl, 2 mM) was added to PBS (10 ml) and this mixture was combined with calcein-AM stock solution (5 µl, 4 mM) to prepare the stain. 500 µl of this reagent was added into each well and incubated for 20 minutes at room temperature. Cell viability was then observed and evaluated using fluorescence microscopy (LEICA Instruments, Milton Keynes, UK) on Image Capture Pro software.

The alamarBlue is a cell metabolic activity assay that can provide cell viability and proliferation assessment. HDF cells were seeded onto the control and discs of PCL, PCL-PNIPAAm, PCL-PNIPAAm-Cur membranes (16 mm diameter, n=4) in 24-well plates at a density of 1 x 10^4^ cells/ml, determined using a haemocytometer and trypan blue staining. 1 ml of media was added in to each well. After incubation of 1, 3, and 7 days under standard cell culture conditions, 100 µl of alamarBlue dye (10% v/v) was added to each well and incubated for 3 hours at 37 °C. Next, 150 µl of the solution was transferred into a 96-well plate and the optical density measurements were performed at 560 nm using a microplate reader (Tecan Infinite M200, Tecan Infinite Ltd., Reading, UK).

### 2.6 Statistical analysis

The experiments were repeated in triplicates unless stated otherwise. All quantitative results are expressed as mean values ± SD. A one-way analysis of variance (ANOVA) with Tukey’s test was used to determine statistical significance. A p-value < 0.05 was considered significant (*). All data was analysed using Microsoft Excel 2019 and presented using Origin Pro 2020. ImageJ software was used to develop the microscopy images.

## 3. Results

### 3.1 Characterisation of PCL, PCL-PNIPAAm and PCL-PNIPAAm-Cur

#### 3.1.1 Lower critical solution temperature (LCST) measurements

LCST transition behaviour was investigated through UV-vis measurement (563nm) in a range of temperature from 4[ to 45[ (Figure 1). The midpoint temperature (32.1□) of between the highest and the lowest absorbance were recorded as the LCST of PNIPAAm. As the temperature of the polymer solution increased, the colour of the solution changed from clear at lower temperatures to cloudy at higher temperatures.

**Figure 1.**
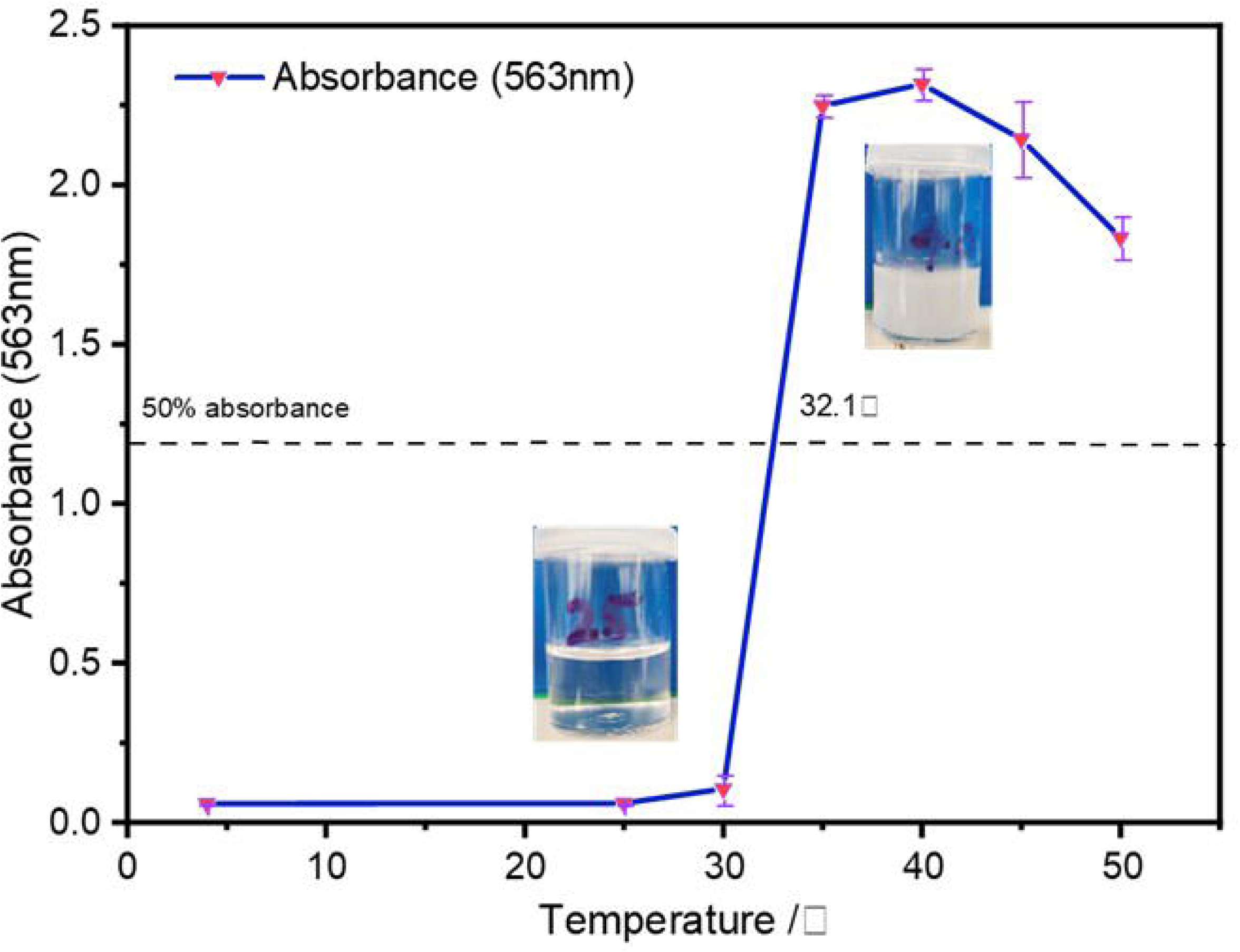
LCST behaviour of PNIPAAm. The value of UV-vis absorbance surged in a temperature range of 30-35[. The LCST is performed at the midpoint of the highest and lowest absorbance.

#### 3.1.2 Chemical composition analysis

Incorporation of PNIPAAm onto PCL nanofibrous surface was confirmed via ATR-FTIR spectroscopy. **Figure 2** shows two ATR-FTIR spectra of increased intensity in the modified PCL samples (PCL-PNIPAAm and PCL-PNIPAAm-Cur) as compared to PCL. A peak at 1647 cm^-1^ indicates the amide I bond, which corresponds to C=O stretching and little C-N stretching of PNIPAAm. Meanwhile, peak at 1565 cm^-1^ corresponds to the amide II bond, arising from N-H stretching and C-N stretching of PNIPAAm. Furthermore, a wider peak between 3550 cm^-1^ and 3200 cm^-1^ was observed for the modified PCL membranes, which correlates to N-H stretching. All samples shown a small sharp peak at 2940 cm^-1^ which is associated with the vibration of aliphatic groups –(CH_2_-)_n_ of PCL. The presence of curcumin can be detected via sharp peaks observed in the PCL-PNIPAAm-Cur spectrum at 1500 cm^-1^ and 1300 cm^-1^ that correspond to C=C bonds and C-O bonds in the phenol groups, respectively (26,28).

**Figure 2.**
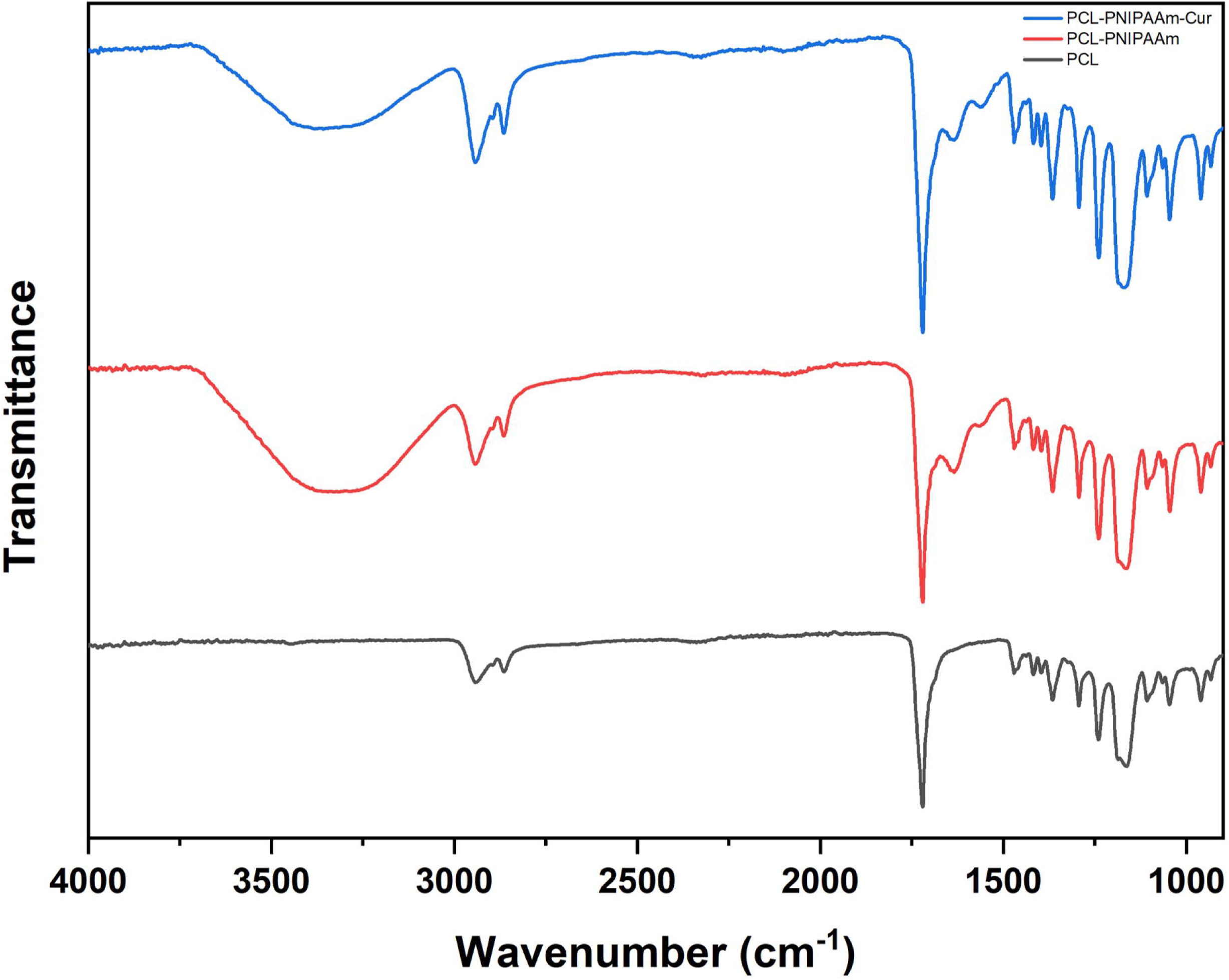
ATR-FTIR spectrum of PCL, PCL-PNIPAAm and PCL-PNIPAAm-Cur membranes; The immobilisation of PNIPAAm onto PCL is confirmed by the presence of both 1647 cm^-1^ and 1565 cm^-1^ peaks, corresponding to the amide I and amide II bonds in PNIPAAm, respectively. These are indicated by the arrows on both the PCL-PNIPAAm and PCL-PNIPAAm-Cur spectra.

#### 3.1.3 Morphology analysis

SEM was employed to study the surface morphologies of PCL, PCL-PNIPAAm, and PCL-PNIPAAm-Cur membranes (**Figure 3**). The PCL sample demonstrated aligned structure of nanofibres at both 1000X and 5000X magnification. However, immobilisation of PNIPAAm on the PCL surface disrupted the structure, as observed in curcumin-loaded PCL-PNIPAAm matrix. Nonetheless, the nanofibre structure was maintained with average diameter 300-800 nm for all three samples.

**Figure 3.**
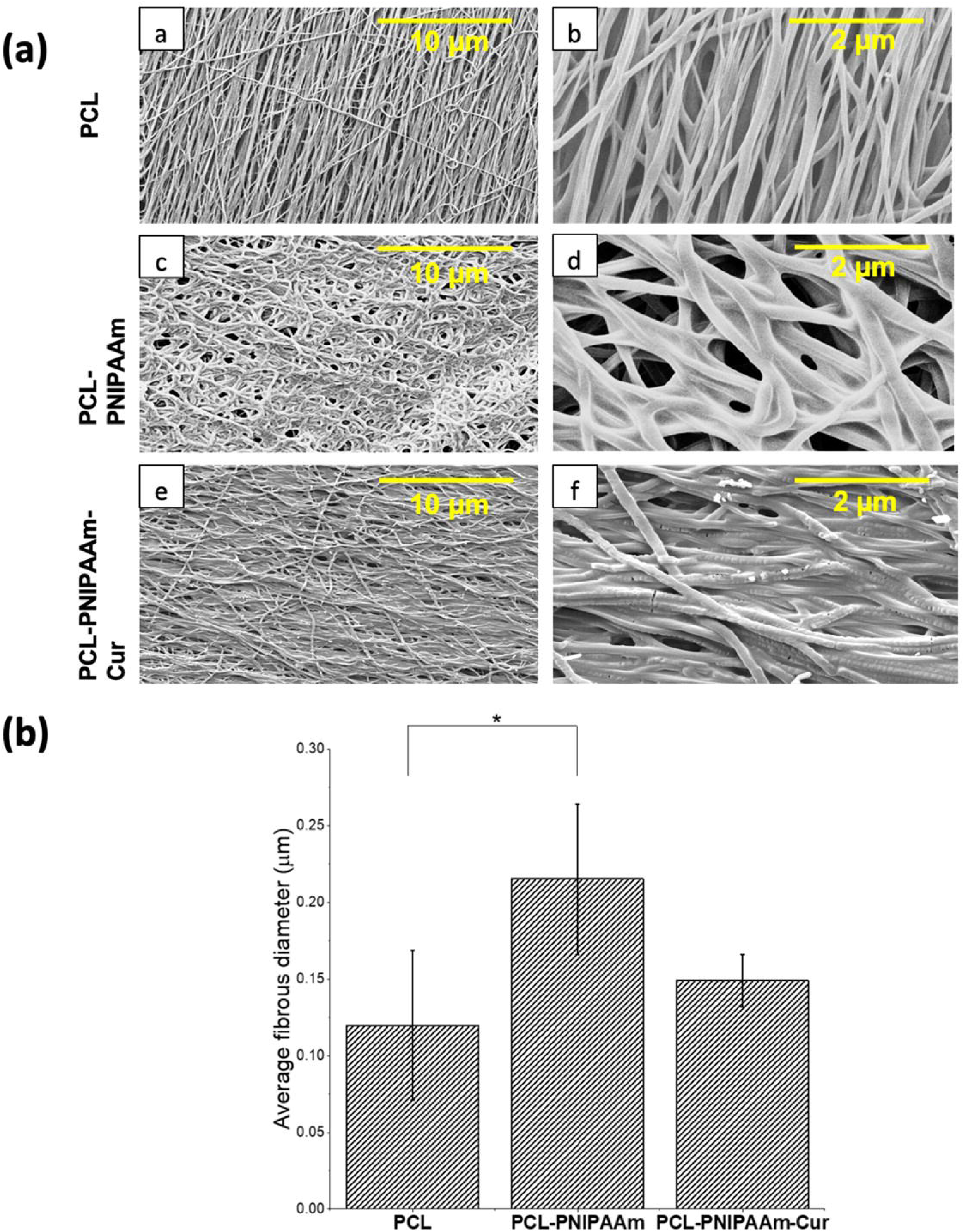
(a) SEM images of the surface morphology and (d) average fibrous diameter of PCL, PCL-PNIPAAm, and PCL-PNIPAAm-Cur membranes; (a-b) The PCL surface exhibits a densely packed and aligned fibrous network. (c-d) This is disrupted in both PCL-PNIPAAm, and (e-f) PCL-PNIPAAm-Cur, although the fibrous structure is maintained. No aggregates of curcumin were observed. Magnification: (a,c,e) 1000X, and (b,d,f) 5000X.

#### 3.1.4 Surface Wettability

Hydrophobicity/hydrophilicity of the PCL, PCL-PNIPAAm, and PCL-PNIPAAm-Cur nanofibrous matrices was determined by measuring the contact angle of water droplets on the surface of each material. The images of the droplets and their corresponding contact angles are shown in **Figure 4**. Water contact angles were averaged over four different samples at room temperature. The average water contact angles for PCL, PCL-PNIPAAm, and PCL-PNIPAAm-Cur were found to be 83 ± 1.2°, 63.5 ± 1.1°, and 68.8 ± 0.9 °, respectively. These results indicate that the hydrophobic nature of PCL was disrupted after conjugation with PNIPAAm and loading with curcumin, resulting in moderate hydrophilicity for both PCL-PNIPAAm and PCL-PNIPAAm-Cur.

**Figure 4.**
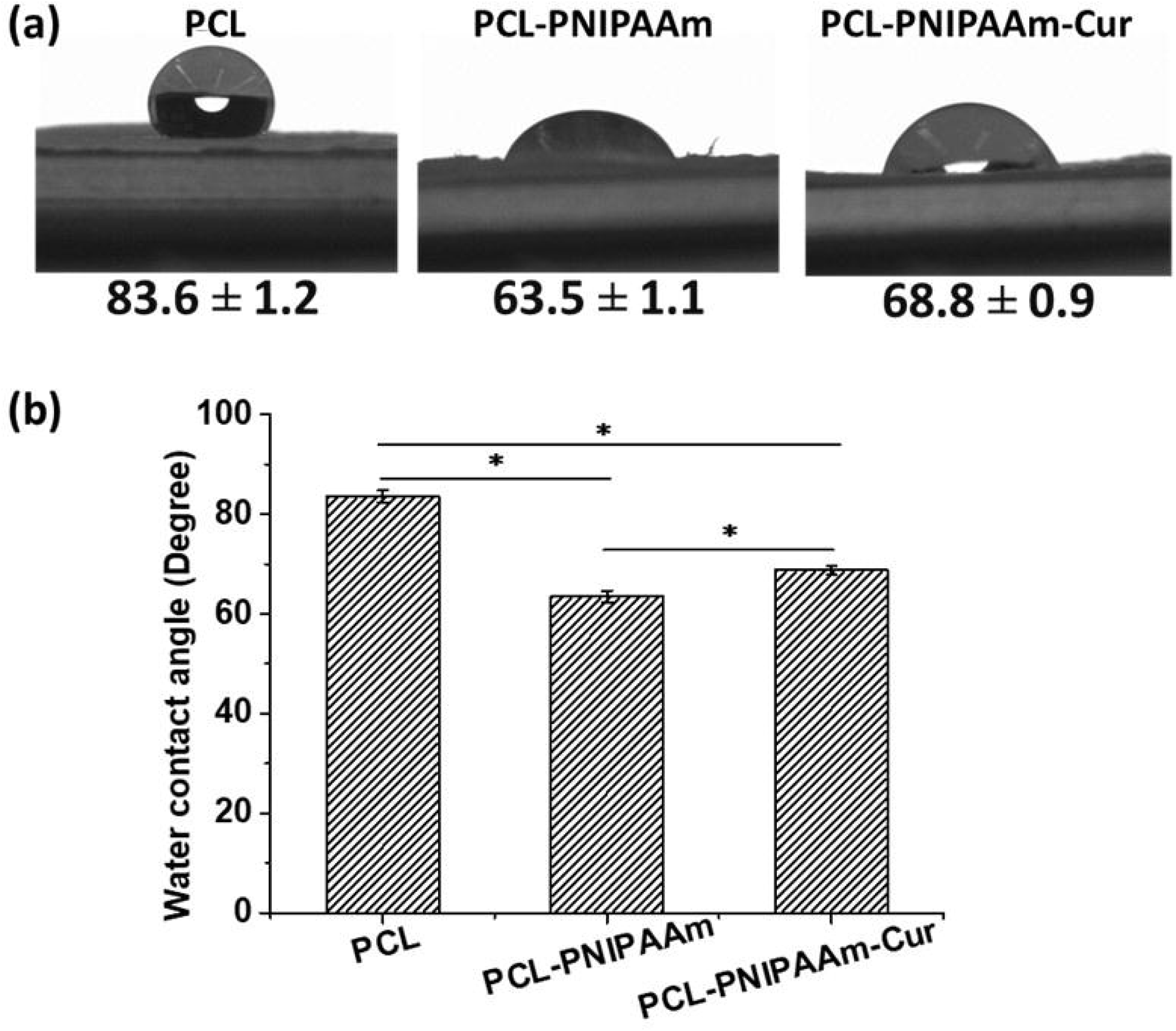
Contact angles of PCL, PCL-PNIPAAm and PCL-PNIPAAm-Cur membranes; (a) Optical image of water drop on nanofibrous membranes and corresponding contact angle, (b) Quantitative values of WCA for PCL, PCL-PNIPAAm, and PCL-PNIPAAm-Cur membranes.

#### 3.1.5 Mechanical testing

The stress-strain curves of PCL and PCL-PNIPAAm were analyzed, shown in **Figure 5** and their mechanical properties were compared. The results, presented in Table 1, showed that the % elongation break and Young’s modulus of PCL-PNIPAAm were significantly higher than those of PCL. However, there was no significant difference in ultimate tensile strength (UTS) between the two samples.

**Figure 5.**
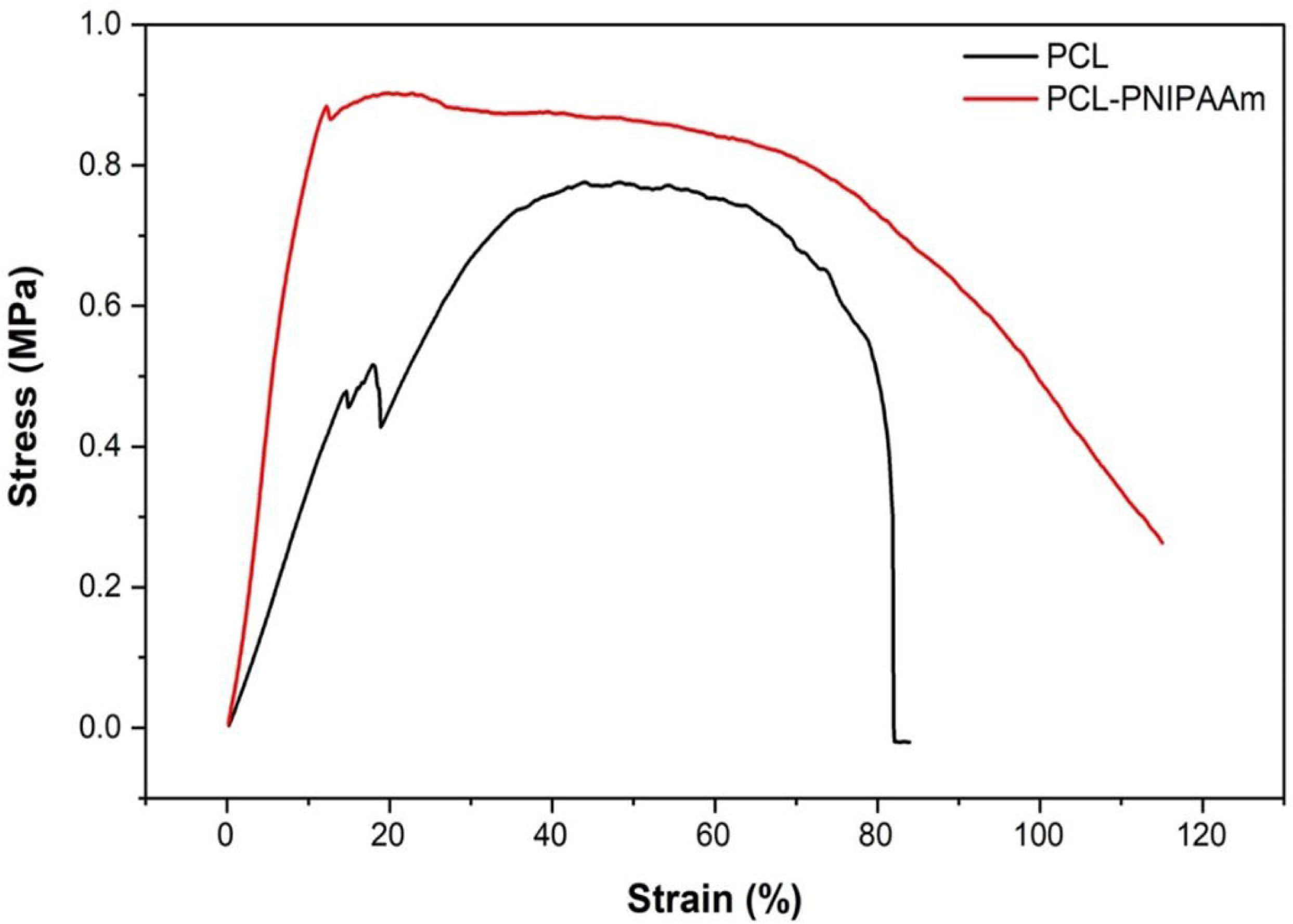
Representative stress-strain curve of both pure PCL and PCL-PNIPAAm membranes.

**Table 1.** The mechanical properties of both PCL and PCL-PNIPAAm membranes. The % elongation and Young’s modulus of the PCL-PNIPAAm membrane was significantly higher than PCL. All values are expressed as mean ± SD for n = 5 (p<0.05).

| Sample | UTS (MPa) | Elongation (%) | Young's modulus (N/m<sup>2</sup>) |
| --- | --- | --- | --- |
| PCL | 0.80 ± 0.10 | 55.00 ± 15.00* | 0.04 ± 1.47 x 10<sup>-4</sup>* |
| PCL-PNIPAAm | 0.90 ± 0.10 | 75.00 ± 20.00* | 0.09 ± 0.001* |

### 3.2 Drug-release profiles

UV absorbance of l_max_ = 432 nm was obtained from curcumin release in media solution and a calibration curve was established at this fixed wavelength to determine the concentration of curcumin released from PCL-Cur and PCL-PNIPAAm-Cur at 4 °C and 37 °C (control). The cumulative release of curcumin against time (1, 2, 3, 4, 5, 24, and 120 hours) was further calculated and the release profiles are depicted in **Figure 6**. After 5 hours i.e., the ‘initial burst’ release profile, 58% and 67% of the drug was released by PCL-Cur at 4 °C and 37 °C, respectively. Additionally, 88% and 58% of the drug was released by PCL-PNIPAAm-Cur at 4 °C and 37 °C, respectively. After 24 hours i.e, the ‘absolute’ release profile, 65% and 74% of the drug was released by PCL-Cur at 4 °C and 37 °C, respectively. Furthermore, 91% and 68% of the drug was released by PCL-PNIPAAm-Cur at 4 °C and 37 °C, respectively. After 120 hours, 71% and 84% of total curcumin was released by PCL-Cur at 4 °C and 37 °C, respectively. Moreover, 94% and 77% of total curcumin was released by PCL-PNIPAAm-Cur at 4 °C and 37 °C, respectively.

**Figure 6.**
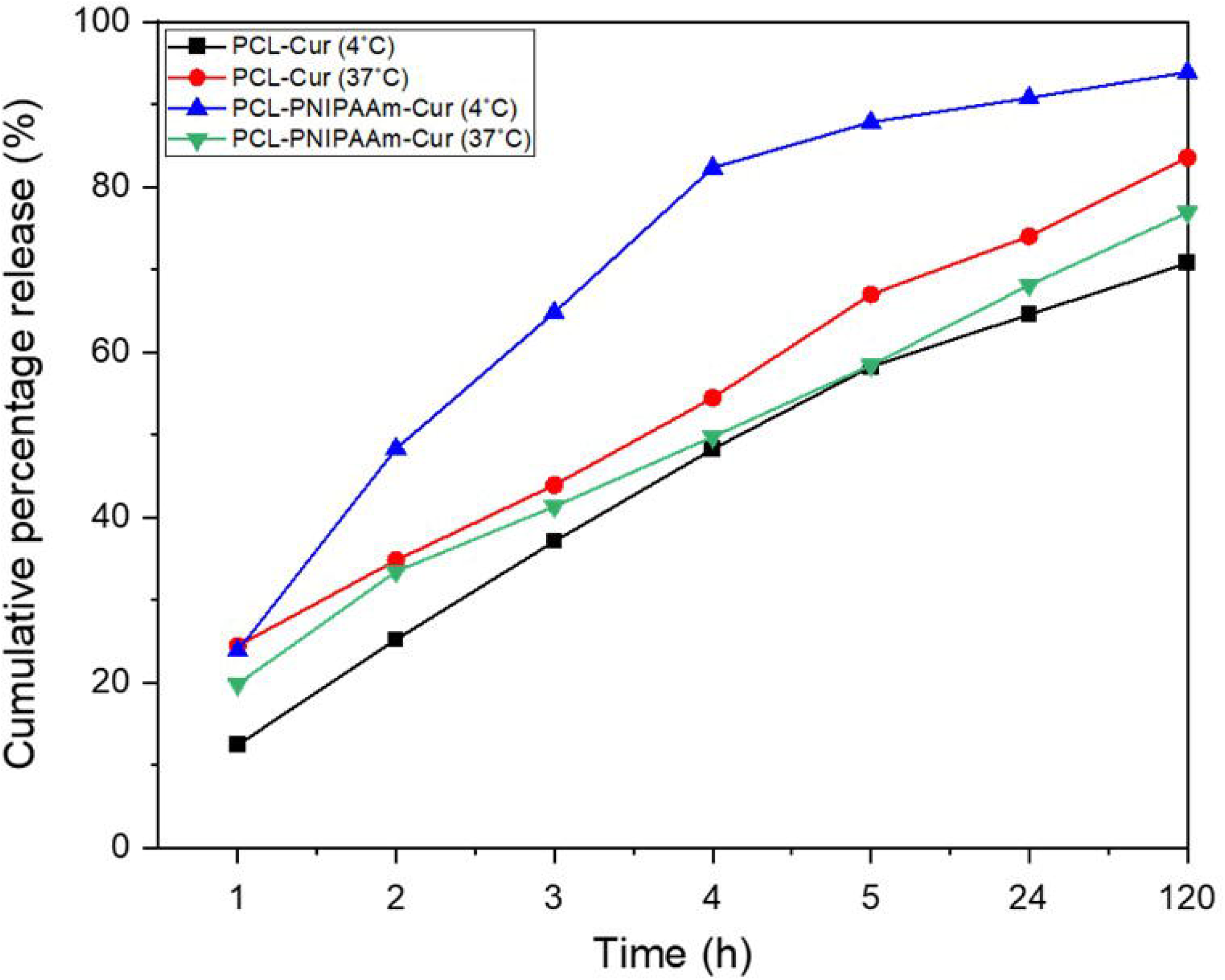
Drug-release profiles of curcumin from PCL-Cur and PCL-PNIPAAm-Cur at 4 °C and 37 °C; The cumulative percentage release of curcumin showed that all membranes exhibited an initial burst release after 5 hours, with PCL-PNIPAAm-Cur exhibiting the highest release (88%) at 4 °C. After 24 and 120 hours, the membranes demonstrated a more steadier release. At 4 °C, PCL-Cur and PCL-PNIPAAm-Cur released 71% and 94% of total curcumin, respectively. At 37 °C, PCL-Cur and PCL-PNIPAAm-Cur released 84% and 77% of total curcumin, respectively.

### 3.3 Cell viability and cell proliferation assessment

To evaluate the cells’ viability and growth against the matrix samples, LIVE/DEAD® Viability/Cytotoxicity assay was used for 7 days duration where cell viability was observed against HDF cells in media (control), PCL, PCL-PNIPAAm and PCL-PNIPAAm-Cur surfaces. LIVE/DEAD images for day 1, 3 and 7 are shown in **Figure 7 (a)**. Auto-fluorescence property of curcumin hinders the imaging, thus no images for PCL-PNIPAAm-Cur sample is included. Although reduced cell viability was observed in both PCL and PCL-PNIPAAm membranes as compared to the control, however very few dead cells were spotted, and the standard spindle-shaped morphology of fibroblast cells was also maintained up to day 7. There were significantly more live cells (green) than dead cells (red) in all scaffolds on days 1, 3, and 7. Reduced cell viability was observed in both PCL and PCL-PNIPAAm compared to the control. Spindle morphology of HDF cells was also maintained in all scaffolds from days 1, 3, and 7. Metabolic activity of HDF cells is assessed via AlamarBlue^TM^ assay in media (control), PCL, PCL-PNIPAAm, and PCL-PNIPAAm-Cur membranes for duration of day 1, 3, and 7. **Figure 7 (b)** shown an increase in absorbance values across all samples where higher metabolic activity recorded for PCL-PNIPAAm and PCL-PINPAAm-Cur scaffolds against HDF cells as compared to pure PCL after day 7.

**Figure 7.**
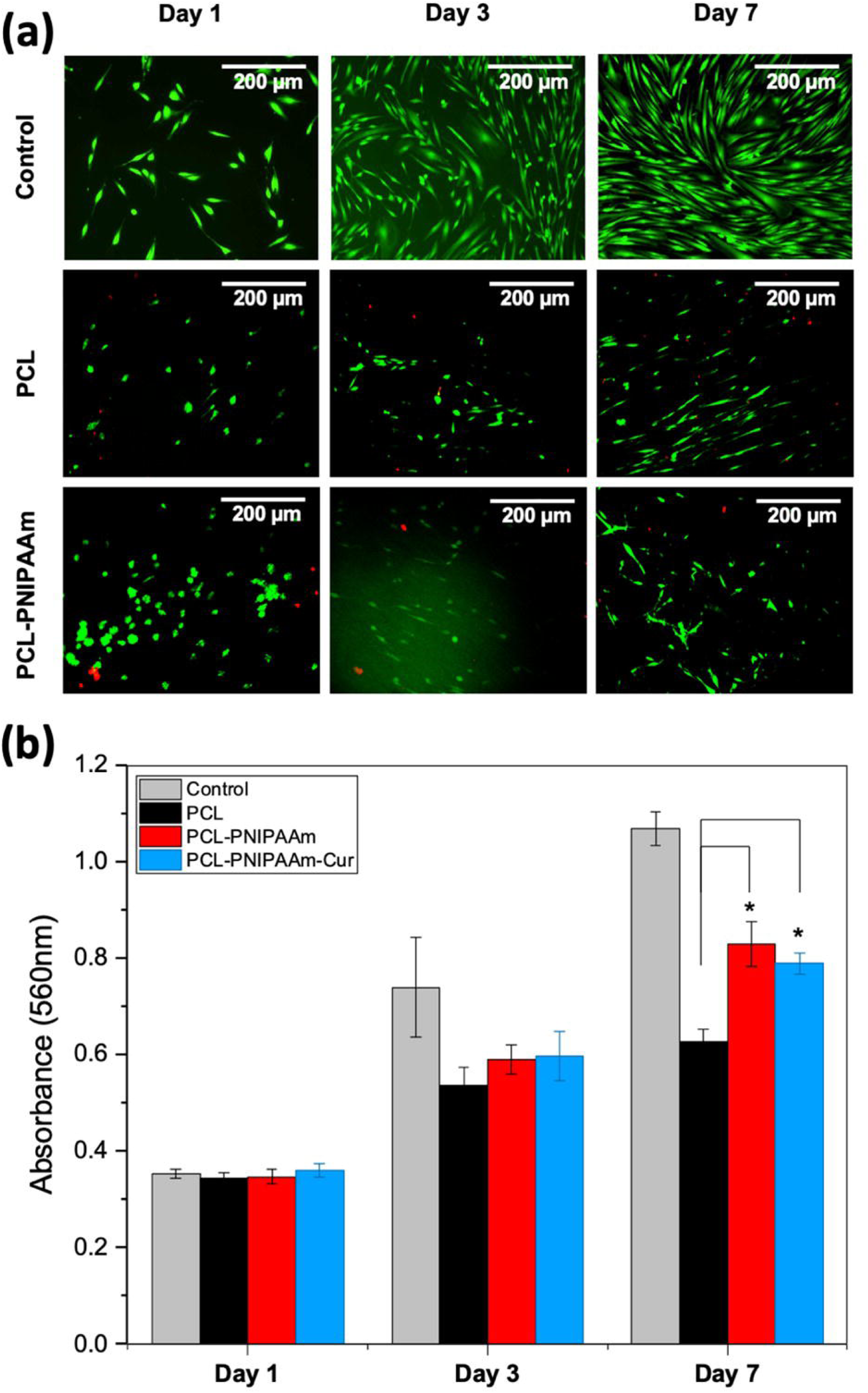
Cellular activity on nanofibrous membranes; (a) Cell viability visualised under a fluorescence microscope using LIVE/DEAD staining on HDF cells seeded in media (control), pure PCL and PCL-PNIPAAm after incubating for 1, 3, and 7 days. **(b)** Metabolic activity of HDF cells seeded in media (control), PCL, PCL-PNIPAAm and PCL-PNIPAAm-Cur membranes observed using alamarBlue staining after incubating for 1,3 and 7 days. Significantly increased proliferation of HDF cells in all scaffolds up to day 7 was observed. The number of HDF cells in PCL-PNIPAAm and PCL-PNIPAAm-Cur was significantly higher than PCL after day 7.

## 4. Discussion

Nanofibrous wound dressings have attracted considerable attention in recent years due to their advantages such as high surface area, porosity, flexibility, and biocompatibility [1]. Among various polymers used for nanofibrous wound dressings, polycaprolactone (PCL) is a promising candidate because of its biodegradability, biocompatibility, and mechanical strength [2]. However, PCL alone has some limitations such as low hydrophilicity, slow degradation rate, and lack of bioactivity [3]. Therefore, PCL is often modified or blended with other materials to enhance its performance for wound healing applications.

One of the strategies to improve the functionality of PCL nanofibers is to conjugate them with poly(N-isopropylacrylamide) (PNIPAAm), which is a well-known thermoresponsive polymer that exhibits a lower critical solution temperature (LCST) around 32°C [4]. Below the LCST, PNIPAAm is hydrophilic and swollen, while above the LCST, it becomes hydrophobic and collapsed. This property allows PNIPAAm to act as a smart drug delivery system that can release drugs in response to temperature changes [5]. For example, when PNIPAAm-conjugated PCL nanofibers are applied to a wound site, they can release drugs at body temperature (37°C) and stop the release at room temperature (25°C). This can prevent drug leakage and enhance the therapeutic efficacy of the wound dressing.

Another strategy to improve the bioactivity of PCL nanofibers is to load them with curcumin, which is a natural polyphenol derived from turmeric that has anti-inflammatory, antioxidant, antibacterial, and wound healing properties [6]. Curcumin can modulate various molecular pathways involved in inflammation, oxidative stress, infection, and tissue regeneration, thus facilitating the wound healing process [7]. However, curcumin has some drawbacks such as low solubility, stability, and bioavailability [8]. Therefore, curcumin needs to be encapsulated or incorporated into suitable carriers such as nanofibers to overcome these challenges and enhance its delivery to the wound site.

In this study, we aimed to develop a novel thermoresponsive nanofibrous scaffold for wound healing applications by grafting PNIPAAm-NH2 onto PCL nanofibers. The grafting process was based on an amidation reaction between the carboxylic acid groups generated by NaOH treatment on the PCL surface and the amine groups of PNIPAAm-NH2. The reaction scheme is illustrated in Fig 8. We confirmed the successful grafting of PNIPAAm-NH2 by FTIR spectroscopy, which showed the characteristic peaks of amide I and amide II bonds at 1647 cm-1 and 1565 cm-1, respectively. The surface modification of PCL enhanced its biological and physicochemical properties, such as hydrophilicity, biodegradability, and bioactivity.

**Figure 8.**
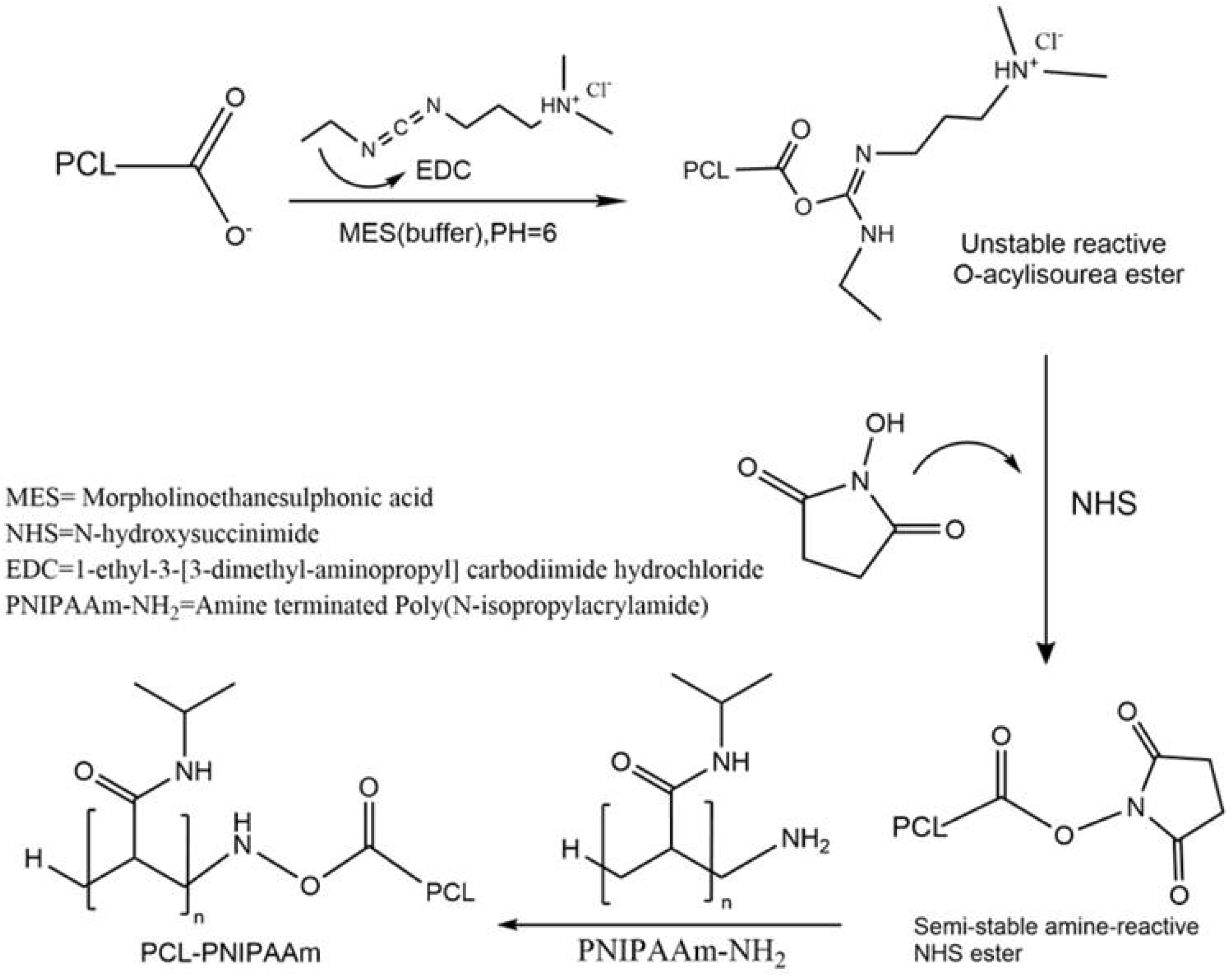
Reaction scheme of an amidation reaction between PCL and amine terminated PNIPAAm to form PCL-PNIPAAm.

We used the “grafting-to” technique, which is a simple and versatile method to conjugate preformed end-functionalised polymers to reactive substrates. In this case, we modified PNIPAAm with an amine group at one end and grafted it to the activated carboxylic acid groups of PCL nanofibers. This resulted in a strong covalent linkage between PCL and PNIPAAm, which imparted thermoresponsive behavior to the nanofibrous scaffold. The grafting density of PNIPAAm was limited by the steric hindrance of the existing PCL layer, which hindered the diffusion of PNIPAAm chains to the microsphere surface. This phenomenon was also observed by Nguyen et al., who reported similar FTIR peaks for PNIPAAm-grafted PCL nanofibers [9]. To further improve the bioactivity of the scaffold, we loaded curcumin, a natural anti-inflammatory and wound healing agent, into the nanofibers. The presence of curcumin was detected by FTIR spectroscopy, which showed the peaks of C=C and C-O bonds in the phenol groups at 1500 cm-1 and 1300 cm-1, respectively, as previously reported by Sahoo et al [10].

The SEM images of PCL, PCL-PNIPAAm and PCL-PNIPAAm-Cur nanofibers are shown in Figure 2. The surface morphology of the nanofibers was not significantly affected by the grafting of PNIPAAm and the loading of curcumin, as they maintained a smooth and bead-free structure. However, some disruption of the alignment was observed in PCL-PNIPAAm and PCL-PNIPAAm-Cur membranes, which could be attributed to the hydrophobic interactions among the three components. The fiber diameter distribution was multimodal, ranging from 300 to 800 nm, and no curcumin aggregates were detected on the surface. The preservation of the nanofibrous structure is important for mimicking and recreating the functionality of the extracellular matrix (ECM), which regulates cell adhesion, differentiation, and proliferation. Moreover, the high surface area of the nanofibers facilitates better endothelial and epithelial cell growth, which are crucial factors for wound healing. Kumar et al. reported that fibers with a diameter ranging between 350 and 1100 nm were optimal for fibroblast matrix deposition and proliferation [11]. Merrell et al. also reported similar results for curcumin-loaded PCL nanofibers, with a diameter ranging from 200 to 800 nm [12].

The water contact angle measurements of PCL, PCL-PNIPAAm and PCL-PNIPAAm-Cur samples are shown in Figure 3. The hydrophilicity of the nanofibers is a crucial property for wound dressings, as it affects the moisture retention, permeability, and biocompatibility of the materials. PCL exhibited poor hydrophilicity, as indicated by its high contact angle of 83° ± 1.2°, due to the presence of CH_3_ groups in its chemical structure. Previous studies recorded higher contact angles of PCL, which some suggested that this may be due to increase in surface roughness [13]. The grafting of PNIPAAm significantly decreased the contact angle to 63.5° ± 1.1°, mainly due to its random coil conformation that leads to better water absorption in its hydrophilic state below LCST (32 °C). The loading of curcumin further decreased the contact angle to 68.8° ± 0.9°, compared to PCL and demonstrating increased hydrophilicity [14]. Although curcumin is insoluble in water, the oxygen in its chemical structure can donate protons to form bonds with water molecules. Moderate hydrophilicity is vital to ensure better haemostasis, inflammation, proliferation, and tissue remodelling of a wound dressing [15–17].

The cytotoxicity and hydrophilicity of the nanofibrous scaffolds are important parameters for evaluating their suitability as wound dressings [18]. Several natural polymers, such as chitosan, alginate, collagen, and gelatin, have been used for wound healing applications due to their low cytotoxicity and high hydrophilicity [19]. However, these polymers also have some drawbacks, such as poor mechanical strength, rapid degradation, and low antimicrobial activity [20]. Therefore, synthetic polymers, such as PCL, have been combined with natural polymers or bioactive agents to improve their performance [21]. In this study, we conjugated PNIPAAm, a thermoresponsive polymer, onto PCL nanofibers and loaded them with curcumin, a natural anti-inflammatory and wound healing agent [5]. We hypothesized that the PNIPAAm-PCL nanofibers would exhibit tunable hydrophilicity and hydrophobicity depending on the temperature, which would enable controlled release of curcumin and prevent bacterial infection [22]. We also expected that the curcumin would enhance the bioactivity and biocompatibility of the nanofibers.

The mechanical properties of PCL and PCL-PNIPAAm nanofibers were analyzed by DMA. There was no significant difference in the ultimate tensile strength (UTS) of PCL (0.80 ± 0.10 MPa) and PCL-PNIPAAm (0.9 ± 0.10 MPa). The UTS values obtained in this study were higher than those reported by Ranjbar-Mohammadi and Bahrami [23] for PCL nanofibers, which may be due to the difference in electrospinning parameters. To the best of our knowledge, there is no previous study reporting the mechanical properties of PCL-PNIPAAm nanofibers, hence more research is needed for better comparison. The elongation at break and Young’s modulus of PCL-PNIPAAm nanofibers were higher than those of PCL nanofibers, indicating improved elasticity and ductility of the PCL-PNIPAAm matrix. Ranjbar-Mohammadi and Bahrami also reported a significant increase in tensile strength and elongation at break of PCL-gum tragacanth nanofibers loaded with curcumin [24]. These results suggest that durability and optimal strength are crucial for an ideal wound dressing to withstand the external environment, dressing changes, and wound contraction.

The in vitro release profile of curcumin from PCL-Cur and PCL-PNIPAAm-Cur nanofibers was studied by UV-Vis spectroscopy. The characteristic peak of curcumin release in media solution was determined at λmax = 432 nm, similar to the value of λmax = 427 nm reported by Sharifisamani et al. for electrospun PEG-PLA-PCL yarns loaded with curcumin [25]. An initial burst release within the first 5 hours was observed from both the PCL-Cur and PCL-PNIPAAm-Cur membranes at 4 °C and 37 °C. The rapid release of curcumin may be due to its high diffusivity in the polymer matrix and its high solubility in the media solution. In addition, the dimensions and uniformity of the fibers may also affect the release kinetics. Bose et al. observed a higher total release of curcumin (39%) with the addition of PCL-PEG on a hydroxyapatite matrix compared to hydroxyapatite alone (16%) after the first 24 hours [26]. They attributed this to the increased hydrophilicity of the drug with the incorporation of PCL-PEG in their study. Similarly, in our study, the contact angle results of PCL (83° ± 1.2°), PCL-PNIPAAm (63.5° ± 1.1°), and PCL-PNIPAAm-Cur (68.8° ± 0.9°) demonstrated significantly increased hydrophilicity upon conjugation of PNIPAAm and subsequent loading of curcumin.

Theoretically, PCL-PNIPAAm-Cur at 37 °C should show no release of curcumin, as this temperature is above the LCST of PNIPAAm (32 °C) and should therefore encapsulate the curcumin in its collapsed hydrophobic globular state [27]. The release may be due to washing out the PCL-PNIPAAm-Cur membranes in between experiments, releasing some curcumin on the outskirts of the nanofibres. Despite this, the 88% release of curcumin by PCL-PNIPAAm-Cur at 4 °C in the initial burst release profile offers promising implications. This is because the membrane can serve as an emergency wound dressing for injuries such as burns, gun and stab wounds to stop blood leaking and provide instant antimicrobial agents, which can be achieved by incorporating and applying the smart dressing with an ice pack and release drugs on demand. This can be self-applied and therefore not burden healthcare clinics, thus also improve patient compliance. However, different concentrations of curcumin release must be studied for future work to enhance evaluation of toxic side effects. Curcumin can downregulate the NF-kB pathway and promote apoptosis in various cells in a dose-dependent manner [28]. Studies have also suggested that high concentrations of curcumin (∼25 µM) can have adverse effects on cell survival (less than 50%) [29].

A >90% percent release of the drug after 24 and 120 hours by PCL-PNIPAAm-Cur at 4 °C is also useful for the assistance of curcumin in the inflammatory, proliferative and tissue remodelling stages for normal wound healing. This can provide maximum therapeutic potential due to curcumin’s strong antioxidant and anti-inflammatory activities [30]. PCL-Cur at 4 °C demonstrated a steadier release than PCL-Cur at 37 °C in both the initial and absolute release profiles, releasing 71% of total curcumin after 120 hours, and could be investigated further as a controlled-release platform. The experiment should also be repeated to improve the reliability of the results, and to obtain an evaluation of any significant difference with the drug-rate release.

Metabolic activity of HDF cells was studied in terms of cytotoxicity and cell growth against all samples. LIVE/DEAD Viability/Cytotoxicity Kit and alamarBlueTM Cell Viability were analysed for samples in media (control), PCL, PCL-PNIPAAm and PCL-PNIPAAm-Cur membranes for duration of days 1, 3, and 7 under standard cell culture conditions. Results shown a significant presence of live cells as compared to dead cells up to day 7, although less numbers observed than control sample. Similar findings by Nguyen et al. which utilised CCK-8 assay kit on PNIPAAm-conjugated PCL membranes also shown more live HDF cells observed for both PCL and PCL-PNIPAAm scaffolds up to day 7 of incubation, with reduced viability as compared to the control [31]. It was also reported higher HDF proliferation rate for PCL-PNIPAAm as compared to PCL, similarly to our findings for alamarBlue assay.

It is very important that our promising cell viability findings could facilitate further research in the future to address the critical role of cells in different physiological stages of wound healing. Cells are recruited to the wound site during the inflammation stage, where chemo-attractants (e.g., PDGF) [32] and inflammatory cytokines such as TNF-α and interleukin-1 beta (IL-β) [33] are released from the provisional matrix consisting of platelets and macrophages. Within 24-48 hours post injury, cells will infiltrate and degrade the fibrin clot through the production of MMPs [34] and synthesise new ECM components, such as glycoproteins, glycosaminoglycans, proteoglycans, thrombospondin, laminin, collagen, hyaluronic acid, fibronectin, and fibrin [35]. The proliferation stage follows 48-120 hours post injury where fibroblasts secrete keratinocyte growth factor (KGF) [36] which assists in re-epithelialisation. Some fibroblasts differentiate into myofibroblasts, which contribute to angiogenesis through the synthesis of VEGF, a pro-angiogenic growth factor [37]. During the tissue remodelling stage, fibroblasts facilitate the degradation of ECM by proteases and replace the initial type III collagen with type I collagen [38], which then assists wound contraction by using the actin bundles on their structure that extend pseudopodia which can bind to adjacent collagen proteins of the ECM and fibronectin [39]. Our initial metabolic activity showed a potential finding which is favourable for fibroblasts to attach, grow and function in the crucial stages of wound healing, with minimal toxic side effects, although more long-term future studies are needed.

## 5. Conclusions

We have successfully developed and characterized a novel nanofibrous matrix consisting of thermo-responsive polymer PNIPAAm conjugated onto PCL nanofibers loaded with curcumin. We have confirmed the presence of PNIPAAm via amidation reaction and the loading of curcumin within PCL nanofibers by chemical analysis, providing an exciting breakthrough. The matrix’s surface hydrophilicity and mechanical strength have improved, with higher elongation and Young’s modulus of PCL-PNIPAAm compared to PCL, which is a significant milestone in facilitating the potential use of this microcarrier for treating acute and low-exudating wounds. The higher release of total curcumin in the initial burst release and absolute release profiles from PCL-PNIPAAm-Cur at 4 °C compared to 37 °C supports the thermo-responsive transition mechanism of PNIPAAm, making it possible to release drugs on demand for treating emergency injuries such as burns, gun, and stab wounds. The self-application aspect of the smart bandage, using an ice pack, can increase patient compliance and reduce the burden of wounds in healthcare clinics, making it an even more promising prospect. The good cell viability with an increment in proliferation of HDF cells observed for PCL-PNIPAAm-Cur compared to pure PCL signifies the matrix’s ability to facilitate wound healing activities. Our current results hold immense potential in the future as a novel dressing, which opens many opportunities for further studies.

## Supporting information

Graphic abstract

## Acknowledgements

This work was supported by grants from the National Research Foundation of Korea [2018K1A4A3A01064257, 2018R1A2B3003446 and 2019R1A6A1A11034536]. The authors would like to thank Nicola Mordan, George Georgiou, and Graham Palmer for their technical support.

