## Supplementary figures and images for "Nanofibrous Wound Dressing with a Smart Drug Delivery System: Poly(N-Isopropylacrylamide)-Conjugated Polycaprolactone Nanofibers Loaded with Curcumin"

### Graphic abstract

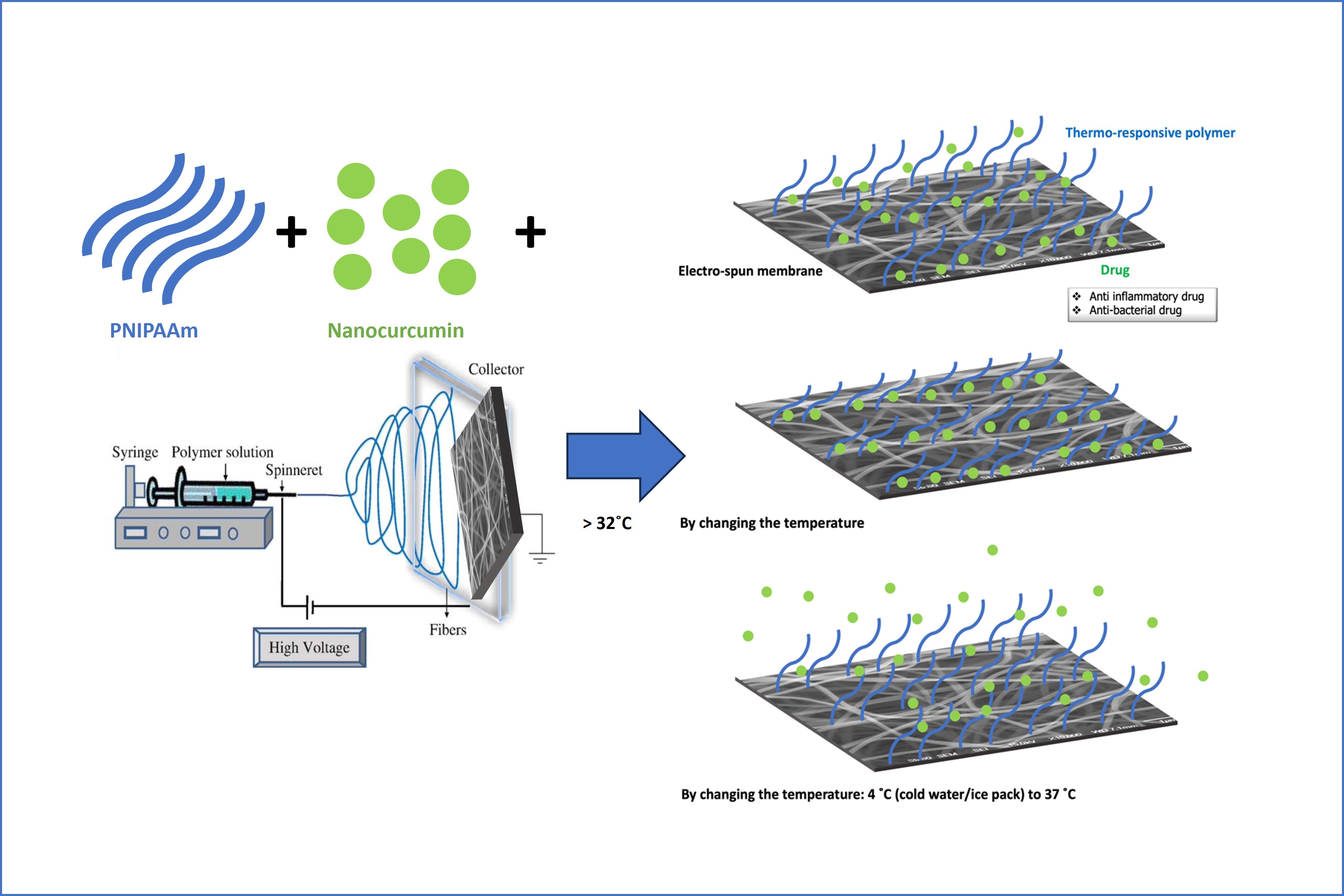
